# pyPhotometry: Open source Python based hardware and software for fiber photometry data acquisition

**DOI:** 10.1101/434225

**Authors:** Thomas Akam, Mark E. Walton

## Abstract

Fiber photometry is the process of recording bulk neural activity by measuring fluorescence changes in activity sensitive indicators (e.g. GCaMP) through an optical fiber. We present a system of open source hardware and software for fiber photometry data acquisition consisting of a compact, low cost, data acquisition board built around the Micropython microcontroller, and a cross platform graphical user interface (GUI) for controlling acquisition and visualising signals. The system can acquire two analog and two digital signals, and control two external LEDs via built in LED drivers. Time-division multiplexed illumination allows independent readout of fluorescence evoked by different excitation wavelengths from a single photoreceiver signal. Validation experiments indicate this approach offers better signal to noise for a given average excitation light intensity than sinusoidally-modulated illumination. pyPhotometry is substantially cheaper than commercial hardware filling the same role, and we anticipate, as an open source and comparatively simple tool, it will be easily adaptable and therefore of broad interest to a wide range of users.

## Introduction

Fiber photometry has emerged as an important tool for behavioural neuroscience, which allows for the measurement of neuronal activity from genetically-defined neuronal populations or axonal projections in superficial or deep brain structures in behaving animals [1–10]. The rapid development of novel genetically encoded fluorescent indicators for calcium [11–15], neurotransmitter release [16–18], and membrane voltage [19–21] suggest that experimental applications for fiber photometry will continue to grow rapidly.

In fiber photometry experiments, excitation light of one or more wavelengths is transmitted to the brain structure of interest through an optic fiber where it excites one or more fluorescent indicators. Light emitted by the indicators is transmitted back through the optic fiber, separated from the illumination light and fluorescence from other indicators using optical filters, and converted to electrical signals by high sensitivity photoreceivers or cameras. These electrical signals are digitised and constitute the data generated by the experiment. Excitation light of different wavelengths may be differentially modulated to allow the fluorescence evoked by each to be demultiplexed from a single photodetector signal, for example to independently measure fluorescence evoked from GCaMP by 470nm and 405nm excitation light. As GCaMP is approximately isosbestic at 405nm (i.e. fluorescence is independent of calcium concentration) this gives a calcium sensitive signal and a calcium insensitive signal that can be used to control for movement artefacts [5].

Turn-key commercial systems exist for fiber photometry data acquisition that are convenient but expensive. Alternatively acquisition can be controlled using generic hardware such as that from National Instruments, but this requires a substantial investment of time to setup as well as proprietary hardware and software licences. We sought to develop an open source acquisition system that offered the convenience of commercial hardware at low cost. As well as democratising access to new experimental methods, open source hardware can help to improve reproducibility as the entire signal acquisition and processing pipeline is open, and facilitate new applications as researchers can modify tools themselves [22–25].

pyPhotometry is a system of hardware and software consisting of an acquisition board and graphical user interface (GUI). The system implements the following functionality: 1) Digitisation of 2 analog voltage signals (at 15-bit resolution) and 2 digital signals. 2) Two constant current LED driver circuits with a 0-100 mA output. 3) Control of data acquisition and online visualisation of signals via the GUI. 4) Streaming of acquired data to disk in a compact binary format. 5) Time-division multiplexed illumination to prevent crosstalk between fluorescence signals and bleed-through of ambient light, with online demultiplexing and visualisation. 6) User documentation at https://pyphotometry.readthedocs.io.

We report the system design and rationale, validation experiments characterising system performance, and data showing its application to recording calcium signals from VTA dopamine neurons.

## Results

### Acquisition board

The acquisition board is built around the Micropython Pyboard, an Arm Cortex microcontroller that is programmed in Python (Figure. 1). Programming the firmware in a high level language allows it to be simple and compact (<200 lines of code), facilitating rapid development. The acquisition board has 2 analog and 2 digital inputs, each a BNC connector. The digital inputs connect directly to general purpose input-output (GPIO) pins on the microcontroller and in principle could be used as outputs, e.g. for triggering closed loop stimulation, though this is not currently supported by the firmware.

**Figure 1.**
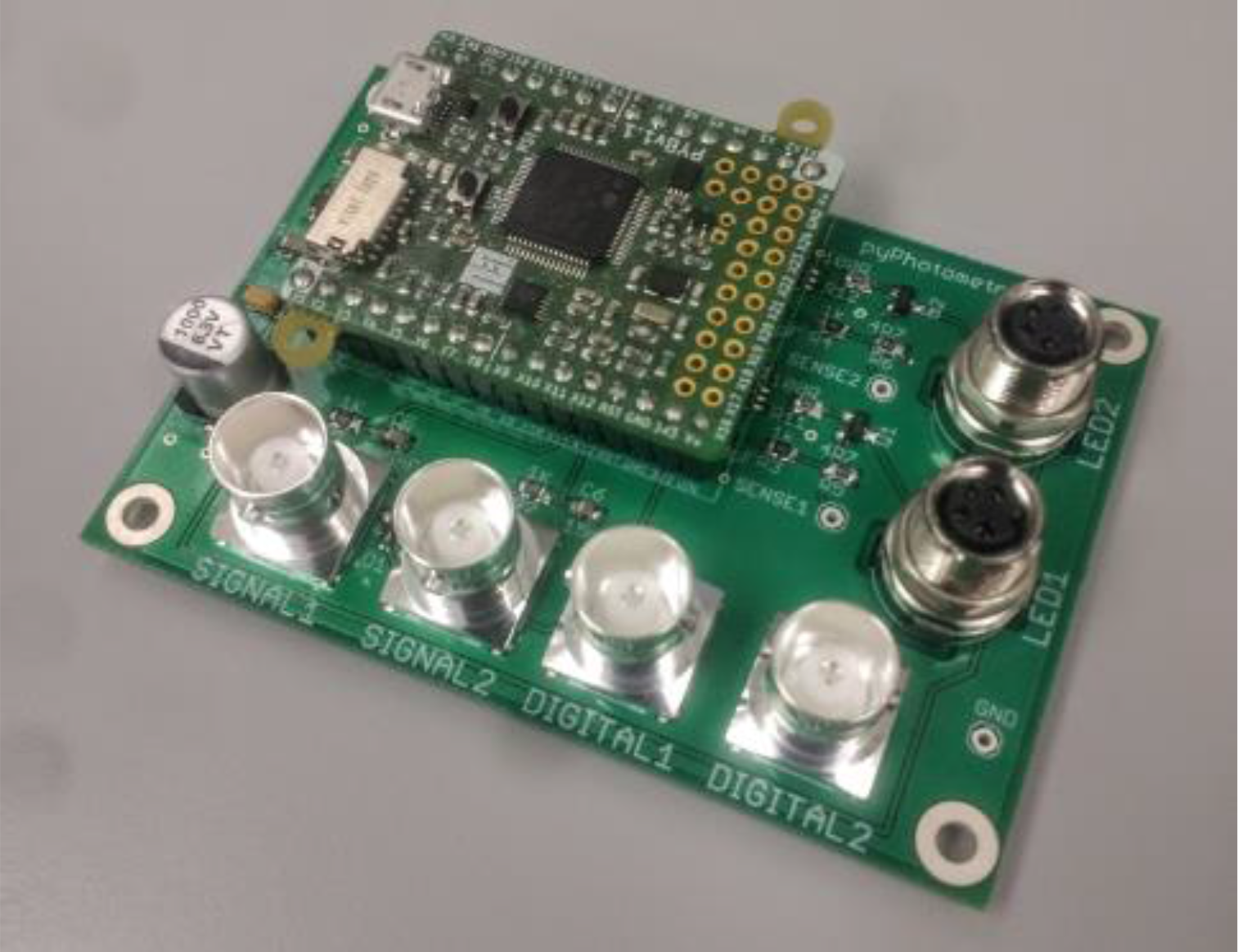
Acquisition board.

Analog signals are acquired using the Pyboard’s analog to digital converters (ADCs). These are 12-bit ADCs with a 0-3.3V range. Oversampling is used to increase the resolution to 15-bit, i.e. for each sample the ADC is read 64 times and the values averaged to give an extra 3 bits of resolution [26,27]. This gives a resolution of ~ 10-4 volts per division of the digitised signal.

The LED driver circuits are voltage controlled current sinks implemented using an op-amp and MOSFET, adapted from Figure 200 of [28]. Their outputs are M8 connectors compatibles with commonly-used connectorised LEDS from Doric Lenses or Thorlabs. The LED driver circuit is linear over a 1-100mA current range and responds to control voltage transients in ~1uS (see *electrical characterisation*).

The acquisition board draws power from the Pyboard’s USB connection which is also used to stream data to the computer. Capacitors on the 5V rail improve transient response and smooth the load presented to the power supply, with a current limiting IC to restrict the inrush current when the board is powered up.

Assembly of the acquisition board requires only standard through-hole and surface-mount soldering techniques. To make a complete photometry system the acquisition board must be paired with LEDs, photoreceivers, filter cubes and other optical components. A complete parts list for the setup we use for green/red two colour experiments (e.g. GCaMP/TdTomato) is provided in the hardware repository. Assembled acquisition boards can be purchased from the Open Ephys Production Site.

### Graphical user interface

The GUI is used to control data acquisition, visualise signals and record data to disk (Figure. 2). It is written in Python and organised into separate modules for communication with the acquisition board, layout of the GUI window, and plotting. The GUI is built using PyQt, a GUI programming toolkit which enables rich GUIs to be implemented compactly-the complete GUI is ~ 600 lines of code. The GUI provides controls for connecting to acquisition boards, setting acquisition parameters, LED currents, the data directory and subject ID, and for starting and stopping data acquisition and recording.

**Figure 2.**
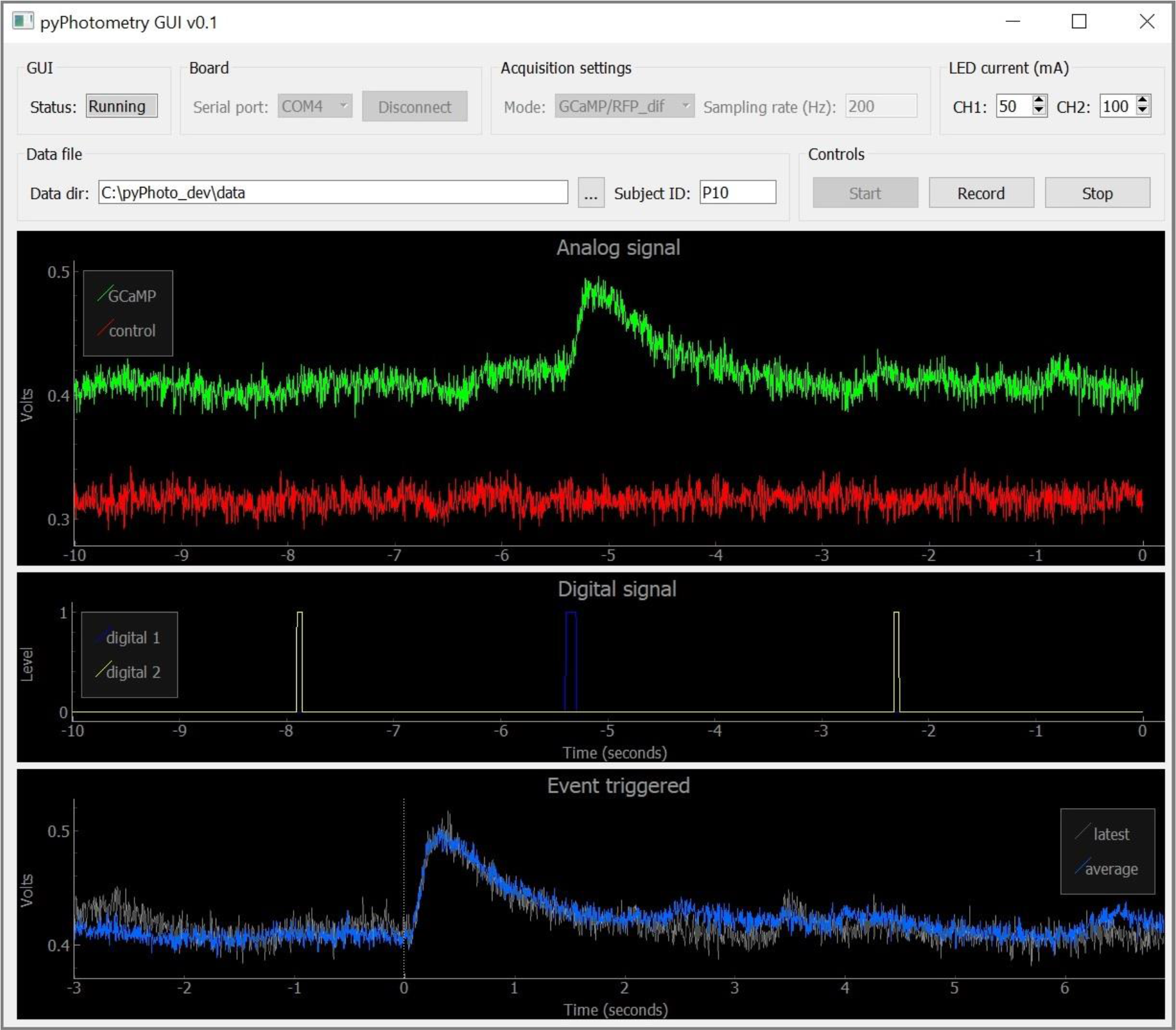
Graphical user interface.

The GUI window has three plots for displaying data implemented using PyQtGraph - a plotting library designed for fast online interactive plotting in GUIs. An *Analog signal* plot displays a scrolling view of the two analog input signals, and is the primary visualisation of the photometry data. A *Digital signal* plot displays a scrolling view of the two digital input signals. An *Event triggered* plot shows a recency weighted event triggered average of analog signal 1 triggered on rising edges of digital signal 1. This allows the average response to an event of interest to be visualised online during data acquisition.

### Data format

pyPhotometry generates binary data files with a *.ppd* file extension. The file format is designed to be straightforward to import into analysis programs while generating files that are no larger than necessary. The user guide provides detailed information on importing *.ppd* files and code for importing *.ppd* files into Python is provided in the *tools* folder of the code repository. Files consist of header information and data. The header is a JSON object that contains the subject ID, start date and time, and acquisition parameters. JSON is a lightweight data interchange format designed to be readable by humans, easy for machines to parse, and is supported by most programming languages [29]. The remainder of the data file contains the analog and digital signals encoded as binary data. Each 2 byte chunk encodes one 15-bit analog signal sample and one digital signal sample, resulting in very compact data files – e.g. a 1 hour recording at 130Hz sampling rate results in a ~1.9MB file.

### Time-division illumination

Photometry experiments often employ sinusoidal modulation of excitation light combined with lock-in amplification - i.e. multiplication of the photoreceiver signal by a sinusoidal reference signal synchronised with the excitation light, followed by low pass filtering [4,5]. This has two advantages over continuous illumination: Firstly, the recovered signal is sensitive only to inputs at the modulation frequency, and thus insensitive to bleed through of ambient light and other noise sources well separated in frequency. As many noise sources have more power at low frequencies, lock in amplification may achieve better signal-to-noise than simply low pass filtering (which also reduces noise bandwidth but does so about 0 Hz) [26]. Secondly, sinusoidal modulation at different frequencies can be applied to different wavelengths of excitation light, and the fluorescence evoked by each independently measured by lock in amplification at the appropriate frequency – a form of frequency-division multiplexing. This is typically used to independently measure the fluorescence evoked in GCaMP by excitation light at 470nm and 405nm.

pyPhotometry uses a different approach to achieve these ends, based on time-division rather than frequency-division principles. Independent readout of fluorescence evoked by different excitation light wavelengths is achieved by alternately switching on LEDs 1 and 2, and acquiring samples of signal 1 when LED 1 is on and of signal 2 when LED 2 is on (Figure 3c). Additionally, baseline subtraction is used to render measurements insensitive to ambient light levels and other low frequency noise sources; for each sample the ADC is read twice; once with both LEDs off to obtain a baseline measurement, and again with the respective LED on. The baseline is subtracted from the sample such that only the difference in light intensity between the LED on and off conditions influences the signal. The acquisition sequence is:

1. Turn LEDs 1 and 2 off, read signal 1 baseline.
2. Turn LED 1 on, read signal 1 sample, subtract baseline and send sample to GUI.
3. Turn LEDs 1 and 2 off, read signal 2 baseline.
4. Turn LED 2 on, read signal 2 sample, subtract baseline and send sample to GUI.

**Figure 3.**
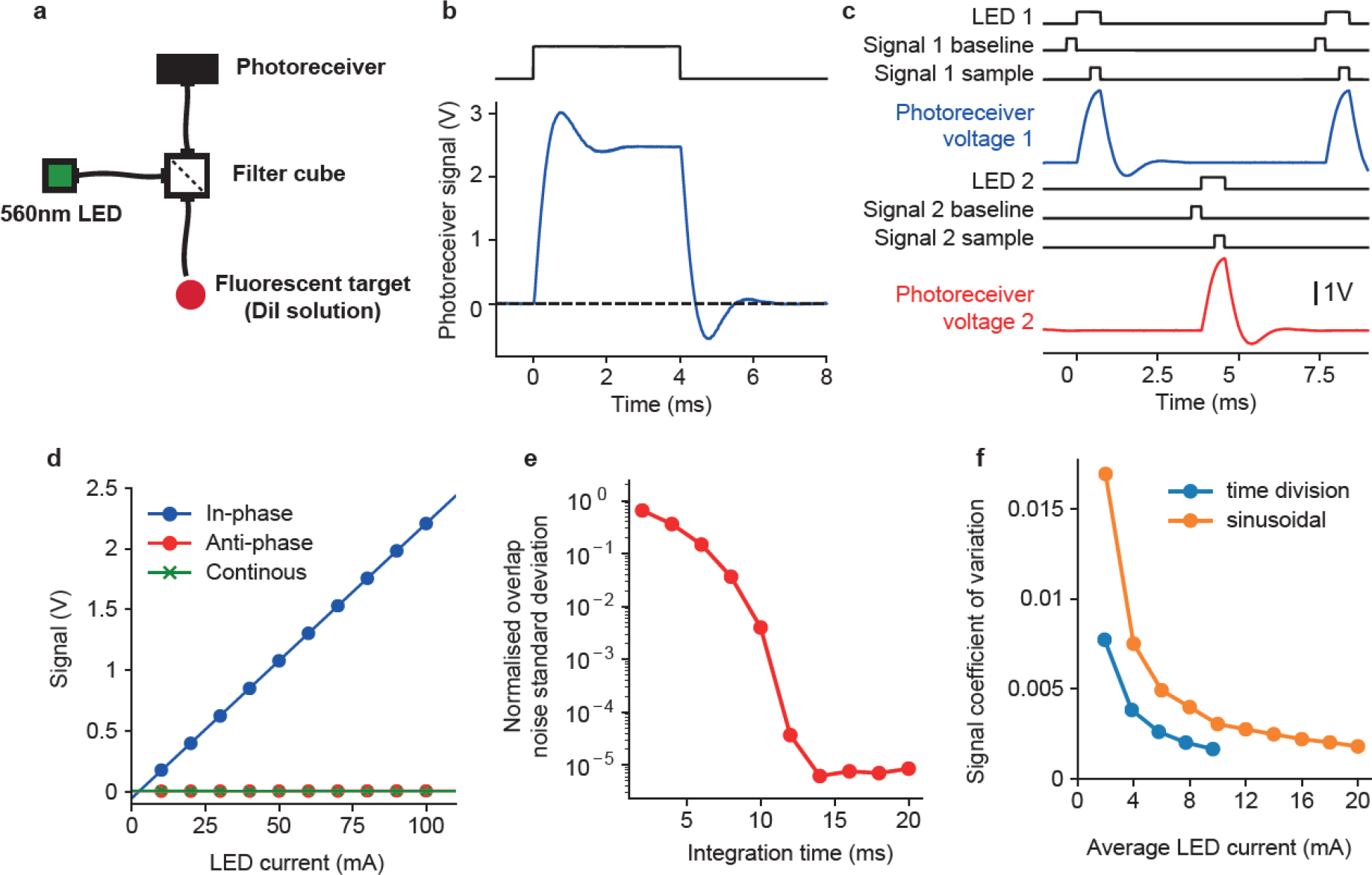
Time-division illumination. **a)**Optical setup for testing time-division illumination and comparison with sinusoidal illumination. **b)**Photoreceiver voltage response to 4ms illumination light pulse. **c)**Timing of events and photoreceiver voltage waveforms for time-division acquisition sequence at 130Hz sampling rate. Black lines show timing of LED illumination and ADC reads of baseline and sample for signals 1 and 2. Blue and red lines show the photoreceiver voltage waveform due to fluorescence evoked by illumination of LEDs 1 and 2. **d)**Baseline subtracted signal as a function of LED current for in-phase illumination, anti-phase illumination and continuous illumination of fluorescent target. **e)**Numerical evaluation of the integration time required for orthogonality between sinusoidal modulations at 211 and 531Hz. Orthogonality was quantified as the standard deviation of the overlap between the two sinusoids normalised by the average overlap of one sinusoid with itself - where overlap between two signals was defined as their product integrated over the time window. **f)**Comparison of noise on signals obtained using sinusoidal illumination with lock-in amplification (orange) and time-division illumination with baseline subtraction (blue). Noise was quantified as the coefficient of variation (standard deviation divided by mean) of the signal, as a function of the average LED current.

This acquisition sequence is used for two acquisition modes, which differ with respect to the analog inputs read to generate signals 1 and 2. The first mode, termed ‘1 *colour time-division’*, uses the same analog input (i.e. a single photoreceiver) for signals 1 and 2. This mode is designed for independently recording the fluorescence evoked at a single emission wavelength by two different excitation wavelengths, for example for measuring the fluorescence from GCaMP due to illumination at 470 and 405nm.

The second mode termed ‘2 *colour time-division’* reads signals 1 and 2 from analog inputs 1 and 2 respectively, i.e. from two separate photoreceivers. This acquisition mode is designed to be used with two fluorophores with different excitation and emission spectra (e.g. GCaMP/tdTomato), allowing separation of both the excitation and emission spectra to be exploited to minimise crosstalk.

### Validation of time-division illumination

To determine the timing of the acquisition sequence for time-division illumination we characterised the time course of photoreceiver voltage signals in response to excitation light transients (Figure 3a, b). The photoreceiver response peaked 0.76ms after the onset of the excitation light and returned to baseline 2.5ms after the light was turned off.

Based on these timings we implemented the acquisition sequence shown in figure 3c. Light pulses from each LED were 0.75ms in duration and occurred at 130Hz, with the two LEDs in anti-phase. Acquisition from the ADCs used 64x oversampling at a 256KHz oversampling rate such that reading a sample took 0.25ms. The baseline for each signal was read immediately before the respective LED was turned on, and the signal sample was read immediately before the LED was turned off.

We assessed the linearity of measurements using this time-division acquisition sequence by measuring the acquired signal as a function of LED current for in-phase illumination of the fluorescent target - i.e. using signal 1 timings for acquisition and LED 1 timings for the light pulses (Figure 3c). In-phase illumination produced a linear relationship between LED current and acquired signal (Figure 3d). We assessed crosstalk between the two signals by measuring the acquired signal as a function LED current for anti-phase illumination – i.e. using signal 1 timings for acquisition but LED 2 timings for the light pulses. The acquired signal was 0 independent of the intensity of out of phase illumination (Figure 3c) indicating no detectable crosstalk. Finally we measured the acquired signal as a function of LED current for continuous illumination – i.e. using signal 1 timings for acquisition with the LED on continuously. Again, the acquired signal was 0 independent of LED current, indicating that baseline subtraction successfully removed sensitivity to steady state light intensity.

We assessed how the time-division illumination used by pyPhotometry compared with the more commonly used sinusoidal modulation in terms of (i) bandwidth (the frequency range of signals that can be measured) and (ii) signal to noise.

The bandwidth achievable using sinusoidal illumination with lock-in amplification is determined by the integration time needed for the modulations of different signals to be orthogonal. Over short integration times the modulations will not be orthogonal and readout will be contaminated by noise due to varying overlap between the target signal’s modulation and the modulation of the other signal. We numerically evaluated the strength of this ‘overlap’ noise as a function of window duration for two signals of equal amplitude modulated at 211 and 531Hz (as used in [5]). Overlap noise decreased smoothly with integration time (Figure 3e) such that the required integration time depended on the noise level deemed acceptable. To achieve an overlap noise standard deviation < 10-3 of the target signal size required a widow duration of ~10ms. Using time division illumination we can acquire samples from two channels at 130Hz, corresponding to a time window of 7.7ms per sample. The time division illumination used by pyPhotometry therefore achieves comparable signal bandwidth to that achieved by the frequency-division methods used in the literature. Both approaches achieve a bandwidth substantially larger than that of GCaMP6f.

Intrinsic noise due to e.g. thermal fluctuations in the photoreceivers, may differentially affect signals acquired using time-division illumination with baseline subtraction and sinusoidal illumination with lock-in amplification. We assessed this experimentally by measuring the coefficient of variation of signal recorded using both approaches from a fluorescent target (DiI solution) as a function of average illumination intensity (Figure 3f). For time-division illumination the pyPhotometry acquisition board was used to acquire signals and control the LED. For sinusoidal illumination a National Instruments USB-6212 BNC board was used to acquire signals and generate the sinusoidal modulation signal, and a Doric Lenses LED driver was used to drive the LED. The optical setup was otherwise identical for the two approaches (see methods). Signals acquired using both methods were low pass filtered at 20Hz to ensure their bandwidth was equivalent.

Noise was substantially lower for signals acquired using time-division illumination, such that a given signal to noise level was achieved at approximately 50% of the average LED current required with sinusoidal illumination. This likely reflects the fact that each LED is on for only 10% of the time during time-division acquisition, such that the illumination light intensity when the signal is measured is 10x higher than the average illumination intensity.

### Electrical characterisation

We performed electrical measurement to characterise the accuracy of the LED driver outputs and the analog inputs.

Linearity of the LED driver was assessed by measuring the LED current as a function of the value written to the Pyboard DAC. LED current was linear in the DAC value across the full range of values (Figure 4a). The standard deviation of LED currents across the 4 driver circuits tested was <1% of the mean current over a range of 1-100mA mean current (Figure 4b). The transient response of the LED driver was assessed by measuring the current waveform in response to a 1ms control voltage pulse (Figure 4c). Rise and fall times of the LED current were of order 1 µS.

**Figure 4.**
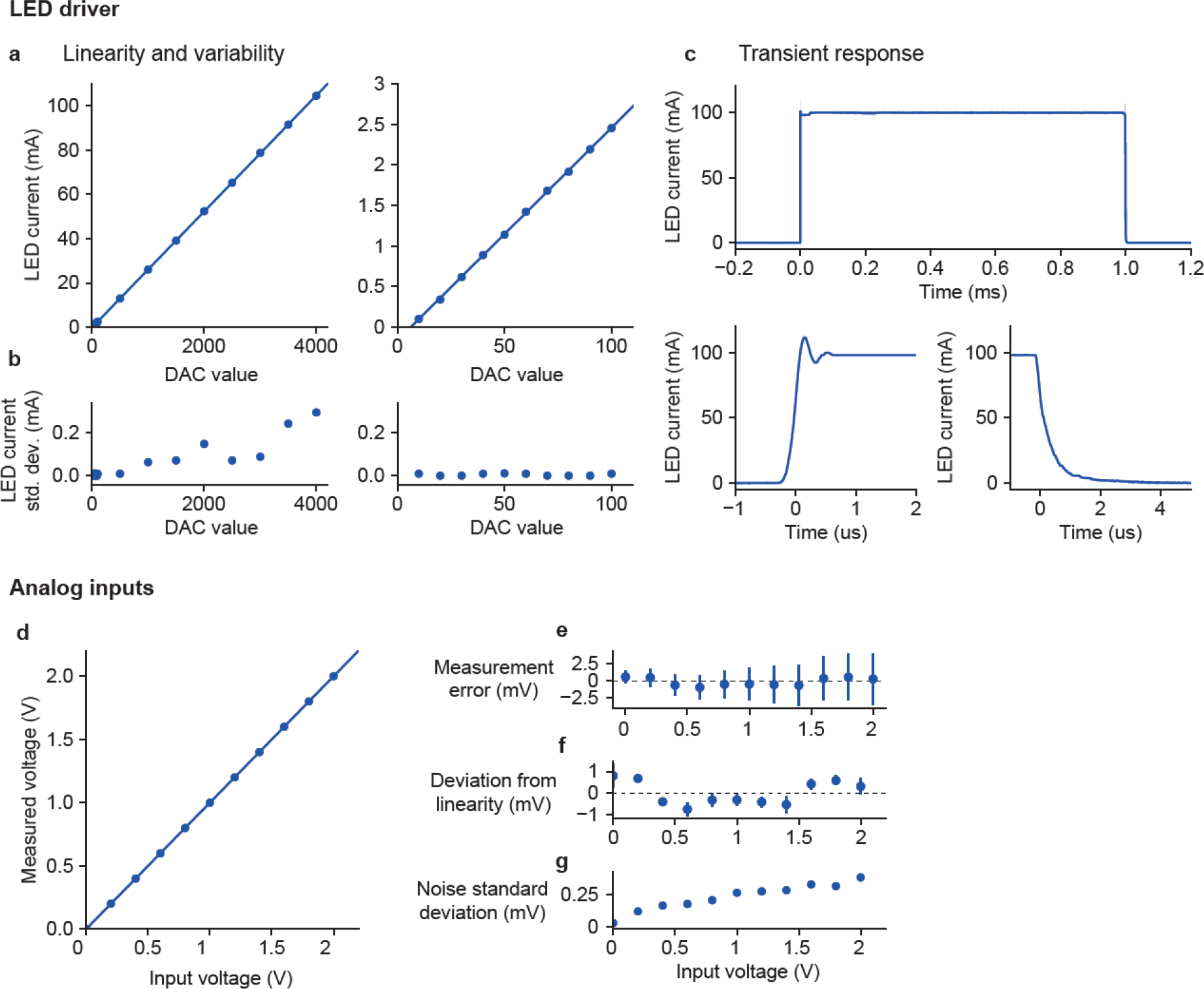
LED driver and analog input characterisation. **a-c)**LED driver **a)**LED current as a function of the value written to the Pyboard DAC in 12 bit mode, points show average measurement across 4 driver circuits tested, lines show linear fit. Left panel – full range of DAC values, right panel – low range of DAC values. The linear fit is the same on both panels. **b)**Standard deviation of LED current across tested driver circuits. **c)**Current waveform in response to 1ms command voltage pulse, top panel - full pulse, bottom panels – rising and falling edges. Line shows average of 32 waveforms, shaded area shows standard deviation (shaded area is hard to see as standard deviation is very small). **d-f)**Analog inputs. **d)**Voltage measured by Pyboard analog inputs as a function of input voltage. Points show average of 8 inputs across 4 Pyboards, line shows linear fit. **e)**Error between measured voltage and input voltage. Points show mean and error bars show standard deviation across inputs. **f)**Deviation from linearity of individual inputs, points show the average residuals from separate linear fits to each input, error bars show the standard deviation of the residuals across inputs. **g)**Standard deviation of noise in the measured voltage, point show the mean across inputs.

To assess the accuracy of voltage measurement using the Pyboard ADCs we used the GUI to read both ADCs at 1KHz while we presented constant voltage inputs. We tested 2 ADCs on each of 4 Pyboards for a total of 8 analog inputs. The measured voltage was very close to linear across the measured voltage range (Figure 4d), with variation in measured voltage across inputs <1% of the mean across the range tested (Figure 4e). We also assessed the extent to which individual inputs deviated from linearity by fitting a least squares linear fit to the measurements from each input separately and plotting the mean and standard deviation of the residuals across inputs (Figure 4f). Deviation from linearity was <1mV across the measured input voltage range and had a characteristic shape across inputs. We measured the noise amplitude of the measured signal. Noise amplitude increased with input voltage but was <0.06% of the measured voltage across the range of voltages tested (Figure 4g).

These results confirm the acquisition board is able to control LED current with high accuracy and temporal precision, and acquire analog signals with good linearity and low noise.

### Dopamine neuron recordings

Having characterised the performance of the hardware we tested its application to recording neuronal activity *in vivo*. Calcium transients were recorded from GCaMP6f expressing VTA dopamine neurons in response to unpredictable reward delivery, which is known to activate a large proportion of these neurons. A water restricted mouse nose-poked in a reward port where 6μL water rewards became available on a random interval 20s schedule. Data was recorded using the *‘2 colour time-division’* acquisition mode to record a GCaMP signal and a movement control signal from co-expressed tdTomato. As can be seen in Figure 5, similar to previous studies [5–7], reward delivery consistently produced a marked, fast transient increase in GCaMP6f fluorescence, while the tdTomato signal was unaffected. This demonstrates the potential utility of using this system for acquiring measures of bulk activity in deep-brain structures in behaving animals. Though we used tdTomato as a movement control channel in these experiments, pyPhotometry can be used for experiments employing 405nm isosbestic illumination of GCaMP as a movement control, using the ‘*1 colour time division’* acquisition mode.

**Figure 5.**
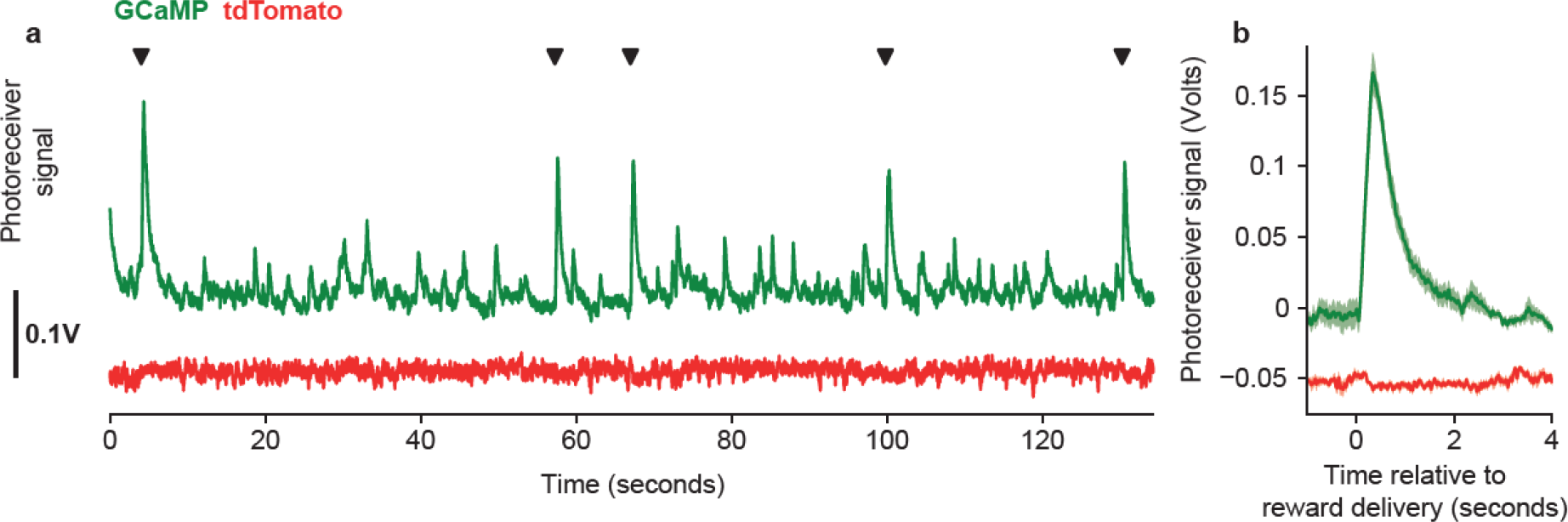
VTA dopamine neuron recordings. **a)**Fluorescent signals acquired with pyPhotometry from GCaMP6f (green) and tdTomato (red) expressed in VTA dopamine neurons. Black triangles indicate times of unpredictable reward delivery. **b)**Response aligned on reward delivery, line shows mean and shaded area shows standard error.

## Discussion

We have developed an open source acquisition board and GUI for fiber photometry data acquisition. The system is compact, convenient, and cheap, and we have characterised its performance in detail.

The system can use time-division illumination to independently measure fluorescence evoked by different excitation wavelengths, combined with baseline subtraction to render measurements insensitive to ambient light. We compared the bandwidth and signal to noise performance of this approach to the more commonly used sinusoidal modulation and lock-in amplification. We found that time-division illumination offered comparable bandwidth to sinusoidal modulation schemes used in the literature, but better signal to noise as a function of illumination light intensity, allowing approximately 50% lower light intensity for a given noise level. Reducing illumination intensity is desirable as it reduces photo-bleaching of fluorophores.

Components of the system lend themselves to adaptation for other applications. The LED driver circuit may be useful in applications where a linear voltage controlled current source with fast transient response is required. The circuit can be modified to handle higher currents by using a MOSFET with higher power dissipation and changing some resistor values. For instance, we use a modified version of the circuit as an analog LED driver for optogenetics.

The GUI code may by useful as a template for applications which require a Python based GUI for data acquisition and plotting. Some pyPhotometry GUI code is shared by pyControl (https://pycontrol.readthedocs.io), a system of open source hardware and software for controlling behavioural neuroscience experiments, also built around the Micropython microcontroller.

While writing this manuscript we became aware of another open source project for fiber photometry called PhotometryBox [30]. This supports generation of sinusoidally modulated control signals (sent to external LED drivers), online demodulation of signals for visualisation (though for analysis demodulation is performed offline), and recording of signals to disk. PhotometryBox uses a microcontroller for generating sinusoidal signals and online demodulation, and a National Instruments board for recording data. The principal differences between the systems are: pyPhotometry has built in LED drivers and is USB bus powered; PhotometryBox must record at higher sampling rates (5KHz) as demodulation is performed offline, resulting in larger data files. pyPhotometry is controlled via a dedicated GUI custom-written for photometry while PhotometryBox uses generic electrophysiology acquisition software (WinEDR) combined with physical controls. Ultimately both systems provide open source photometry data acquisition at a fraction of the cost of commercial systems. We expect this proliferation of open source tools for photometry to both reduce research costs and facilitate the development of novel applications for photometry.

## Methods

### Open source repositories

Design files for the system are at https://bitbucket.org/takam/pyphotometry and https://bitbucket.org/takam/pyphotometry_hardware. User documentation is at https://pyphotometry.readthedocs.io.

### Electrical calibration experiments

A Picoscope 2204A USB oscilloscope was used for all electrical measurements and as a signal generator for testing the analog inputs.

LED current was measured via the voltage across the 4.7 ohm current sense resistors that form part of the driver circuit, using the provided connection points on the board. LED current was measured at 8 DAC values spaced from 500 to 4000 (the DAC takes values from 0 to 4095 in 12 bit mode), and a further 10 values spaced from 10 to 100 to cover the low current range. A single linear fit was made to the full set of points.

### Time-division illumination

Experiments characterising the time-division illumination mode and comparing it with sinusoidally modulated illumination used a Doric Lenses CLED 560nM LED for illumination, a Doric Lenses FCM5 minicube to separate excitation and emission light (excitation filter 555-570nM, emission filter 580-680nM), and a Newport 2151 photoreceiver in DC coupled mode to detect the emitted light. The fluorescent target was a solution of DiI in ethanol, shielded from ambient light and coupled to the minicube via 200um core 0.48NA optical fiber.

To evaluate signal to noise for time-division illumination (Figure 3f) we used the pyPhotometry acquisition board and GUI to both control the LED and acquire signal. For sinusoidal illumination the LED was controlled using a Doric Lenses LEDD_2 driver in low power analog modulation mode. The sinusoidal voltage signal used to modulate the LED was generated by a National Instruments USB-6212-BNC board controlled by WinEDR software (John Dempster, University of Strathclyde). The modulation amplitude was equal to the average value, i.e. the modulation depth was 100%. The USB-6212 was used to record both the LED modulation signal and the photoreceiver signal at 10KHz. Lock in amplification was performed offline using the following steps: 1) Bandpass filtering the recorded modulation and photoreceiver signals around the modulation frequency. 2) Applying a time lag to the modulation signal to phase align it with the photoreceiver signal. 3) Multiplying the lagged modulation with the photoreceiver signal. 4) Low pass filtering the product at 20Hz.

All filtering steps used 4^th^ order zero phase filters, implemented by filtering in the forward and reverse directions using a 2^nd^ order Butterworth filter. The same 20Hz low pass filtering was applied to signals acquired using time-division illumination to ensure noise comparison was made on signals of equivalent bandwidth.

Figure 3e used numerical simulation to assess the bandwidth achievable with the sinusoidally modulated illumination and lock-in amplification used in [5]. Consider using lock-in amplification to measure the amplitude of sinusoidal signal *a(t)* in the presence of sinusoidal signal *b(t)* at a different frequency. Lock-in amplification works by multiplying the input with a sinusoidal reference signal synchronised to the target signal, followed by low pass filtering. The lock-in output at time *T* is:

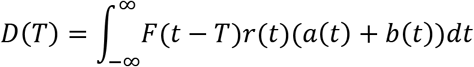

Where *r(t)* is the sinusoidal reference signal and *F*(*t* − *T*) is the impulse response of the low pass filter.

The quantity we plot in figure 3e which we term *normalised overlap noise* is:

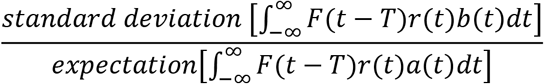

Where the expectation and standard deviation are over measurement time-point *T*. The numerator is the standard deviation of noise on the lock-in output due to overlap between the sinusoidal input *b(t)* and the reference signal *r(t)*. The denominator is the expected contribution of the target signal *a(t)* to the lock-in output.

We evaluated this quantity numerically for sinusoids of equal amplitude at 211 and 531Hz. A Dolph-Chebyshev window with 100dB side lobe attenuation was used as the filter impulse response function *F*(*t* − *T*). We report the overlap noise as a function of the width of this integration window. This window function was chosen because it performed best out of all those tested.

### Dopamine recordings

GCaMP and tdTomato were expressed in VTA dopamine neurons using *AAV1-Syn-Flex-GCaMP6f-WPRE-SV40* (titer 6.2 × 10^13^) and *AAV1-CAG-Flex-tdTomato-WPRE-bGH* (titer 3.1 × 10^13^) viruses (Penn Vector Core) in male *B6.SJL-Slc6a3tm1.1(cre)Bkmn/J* mice. The viruses were mixed and diluted in a ratio of 20% GCaMP6f, 10% TdTomato, 70% saline. Mice were anaesthetised with isoflurane (3% induction, 0.5-1% maintenance), treated with buprenorphine (0.1 mg/kg) and meloxicam (5mg/kg), and placed in a stereotactic frame. The skull was exposed and holes drilled to allow 500nL per hemisphere of the diluted virus to be injected at 1nL/second at AP: -3.3, ML: ±0.4, DV: -4.5mm relative to bregma. Recordings were made through a 200um 0.53NA fiber optic cannula implanted at AP: -3.3, ML: +0.4, DV: -4.3mm relative to bregma. Mice were given additional doses of meloxicam each day for 3 days after surgery, and were monitored carefully for 7 days post-surgery. Prior to recording, mice were put on a water restriction schedule where on training days they received 0.5-1.5mL water from rewards received in the task and on non-training days 1 hour of unrestricted access in their home cage. Mice maintained a typical body weight of >90% pre-restriction levels. Experiments were carried out in accordance with the Oxford University animal use guidelines and performed under UK Home Office Project Licence P6F11BC25. Signals were acquired using the two colour time-division acquisition mode using 470 and 560nm wavelength LEDs respectively for the GCaMP and tdTomato excitation light and 500-540 and 600-680nm emission filters for the GCaMP and tdTomato signals. Acquired signals were bandpass filtered between 0.01 and 20Hz using a fourth order zero phase filter. The full set of optical components used is listed in the hardware repository.

## Author contributions

T.A. designed and built the hardware and software, designed experiments, analysed data and wrote the manuscript. M.E.W. designed experiments and edited the manuscript.

## Acknowledgements

We thank Laura Grima for providing the dopamine neuron recording shown in Figure 5, and for comments on the manuscript. The work was funded by a Wellcome Senior Research Fellowship to Mark Walton (202831/Z/16/Z).

## Data availability

All data and analysis code used in the study are in the file ‘manuscript code and data.zip’ in the downloads section of the code repository: https://bitbucket.org/takam/pyphotometry/downloads/.

